# ExoSloNano: Multi-Modal Nanogold Tags for identification of Macromolecules in Live Cells & Cryo-Electron Tomograms

**DOI:** 10.1101/2024.10.12.617288

**Authors:** Lindsey N. Young, Alice Sherrard, Huabin Zhou, Farhaz Shaikh, Joshua Hutchings, Margot Riggi, Michael K. Rosen, Antonio J. Giraldez, Elizabeth Villa

## Abstract

In situ cryo-Electron Microscopy (cryo-EM) enables the direct interrogation of structure-function relationships by resolving macromolecular structures in their native cellular environment. Tremendous progress in sample preparation, imaging and data processing over the past decade has contributed to the identification and determination of large biomolecular complexes. However, the majority of proteins are of a size that still eludes identification in cellular cryo-EM data, and most proteins exist in low copy numbers. Therefore, novel tools are needed for cryo-EM to identify the vast majority of macromolecules across multiple size scales (from microns to nanometers). Here, we introduce and validate novel nanogold probes that enable the detection of specific proteins using cryo-ET (cryo-Electron Tomography) and resin-embedded correlated light and electron microscopy (CLEM). We demonstrate that these nanogold probes can be introduced into live cells, in a manner that preserves intact molecular networks and cell viability. We use this system to identify both cytoplasmic and nuclear proteins by room temperature EM, and resolve associated structures by cryo-ET. We further employ gold particles of different sizes to enable future multiplexed labeling and structural analysis. By providing high efficiency protein labeling in live cells and molecular specificity within cryo-ET tomograms, we establish a broadly enabling tool that significantly expands the proteome available to electron microscopy.

## Introduction

Cellular cryo-Electron Tomography (cryo-ET) is a powerful method to study the structure and interactome of macromolecules directly in the cell. To access the cell interior at near-native conditions, cells are vitrified and micromachined using Focused Ion Beam (FIB) milling to generate 80-250 nm thick lamellae. The lamella is imaged by transmission electron microscopy (TEM) at different tilt angles to produce a tilt series, which is then aligned. By back-projection one can then generate a three-dimensional tomogram at high spatial resolution that directly represents the structures of intact molecular networks. Recent reviews on the cryo-FIB-ET workflow include [1–4].

Cellular cryo-ET allows structural biology and cell biology to be performed within the same experiment by leveraging preserved structural information in a cellular context. Cellular EM has proven valuable to bridge structural and cellular biology by resolving protein structures inside the cell. So far, complex macromolecular assemblies such as nuclear pore complexes, proteasomes, ribosomes, have been resolved in situ, or their native environment, in various cell lines and conditions. These assemblies represent a small class of macromolecular complexes that are large enough to be unambiguously identified in tomograms. However, approximately 10,000 different proteins were identified by MS/MS to be present in cells at any given time [5]. Specific macromolecular complexes of interest are possible to locate and identify using molecular tags, fluorescence microscopy, and image recognition algorithms. But no method is applicable to image a broad range of proteins embedded in the molecular networks in which they operate. Here, we present a new method, ExoSloNano, to expand the proteome that is accessible to cryo-ET.

### Current labeling technique in cryo-ET

One of the major benefits of cryo-ET, that the sample is vitrified and directly visualized without stain [6], prevents introduction of labels through standard fixation and permeabilization approaches, and is incompatible with powerful oxidative probes such as APEX and APEX2, minisog, ChromEMT [7–9]. Antibodies have been a common choice for immunolabeling in TEM, but they cannot be introduced without fixation and permeabilization, are large, and have limited nuclear delivery. Moreover, a growing consensus argues that poorly vetted commercial antibodies contribute to the reproducibility crisis in basic science [10,11]. Currently available labeling strategies in cellular cryo-EM utilize large macromolecules ranging from 20-70 nm. While such labels are beneficial for visual identification, they may disrupt complex molecular networks within the crowded cellular environment, or affect their formation [12–14].

### Correlative light and electron microscopy (CLEM)

Wide field cryo-fluorescence and super-resolution microscopy (cryo-SRM) can be used for correlated target identification of cells or regions of cells on an EM grid or molecules within a cryo-tomogram [15], with cryo-SRM being actively developed [15,16]. A major challenge with widefield cryo-CLEM is that the resulting fluorescence signal cannot always be overlaid with the field of view of the tomogram. Essentially, the field of view in a cryo-electron tomogram can be as small as 500 nm, while the precision of the fluorescence signal is typically limited to ∼350 nm due to limitations in the optical setup at cryogenic conditions. Thus, implementation remains challenging as diffuse fluorescence puncta can guide the general location within the cell for FIB milling or tilt series acquisition, but not resolve discrete macromolecules [17]. Advances in these areas will continue to provide higher-precision and accuracy in localization of macromolecules, but we expect that a purely fluorescence-based tagging system to correlate single molecules with cryo-ET data will have limitations. Advances in cryo-fluorescence and cryo-Super Resolution Fluorescence microscopy will benefit substantially from labels that can be directly visualized in cryo-ET data in order to find and label molecules ranging from cells to molecules (i.e. tens of microns to the angstrom scale, 5 orders of magnitude).

### Identifying molecular targets in silico

It would be ideal to complement experimental methods to identify novel targets with the growing computational and artificial intelligence programs for particle identification. Template matching (TM) programs include 2D-Template Matching [18], a computationally exhaustive search of highresolution information from a 3-D reference to match to the high-frequency information acquired on a close-to-focus 2-D micrograph. A similar method has been developed for 3-D, though acquired at standard defocus values for cellular cryo-ET [19]. As of yet, the matching of high resolution signatures (2D template matching) is still yet restricted to molecules of sufficient molecular weight [18,20]. Machine learning (ML) programs for particle identification include TomoTwin and DeepFinder [21,22].

The potential of ML for image pattern recognition is undoubtable. However, currently, most ML algorithms act as robust particle pickers-identifying molecules that could be identified by eye, but in an automated way. The added challenge is insufficient priors or ground-truth knowledge with which to evaluate TM and ML outcomes for novel targets. Template matching, machine learning and artificial intelligence are image classifiers, and the low SNR ratio of cryo-ET limits their application to the entire proteome. Finally, there are homologous macromolecules inside the cell that possess high structural similarity to each other. It would be beneficial to have tools to differentiate between these in cryo-ET data.

Small differences in structural features are difficult to separate when matching high resolution features as with template matching procedures when structural differences between such homologous structures are <1 Å^*2*^. For instance, a core nucleosome and nucleosomes containing histone variants have an Root-Mean-Square-Deviation (RMSD) <1 Å^*2*^ yet elicit different functions within the nucleus.

### Gold is king

When imaging vitrified biological material by TEM, the pixel intensity is roughly proportional to the mass of the object [23]. Biological samples contain macromolecules composed of elements of low and similar atomic number (carbon, oxygen, nitrogen, phosphorus), which leads to low contrast in imaging. In contrast, gold nanoparticles provide useful high-contrast markers because of gold’s high atomic number and the condensed structure of the nanoparticles. Immunogold labels from colloidal gold have been used extensively in resin-embedded TEM [24]. However, colloidal gold is large, polydisperse and relies on non-specific interactions for binding [25]. In contrast, nanogold particles are monodisperse, and nanogold labeling of macromolecules has been demonstrated in biochemical systems [26].

Recent work using nanogold as fiducials in cryoconditions include external nangold on the cell surface [27] and endocytosed 2.2 nm gold nanogold particles (Au NP) [28]. Similarly, nanogold-mediated labeling of macromolecules in live cells required endocytosis of the Au NPs [29–31], which may not afford access to all subcellular compartments. However, fully generalizable nanogold labeling for cytosolic and nuclear targets has not yet been demonstrated, and would be a powerful tool for the cellular EM field.

We sought to utilize two modalities-room temperature TEM and cryo-EM in order to leverage the scale and statistical power of the former with the high magnification and details of the latter. This method should be broadly useful to room temperature EM and cryo-ET, and it was developed with challenging targets in mind. Challenging targets would include macromolecules that are <200 kDa, have high structure similarity to other targets, and have a specific suborganellar localization. Finally, TEM labeling within the nucleus has been challenging. The methods presented here would complement the progress within the chromatin organization field and provide more tools for studying chromatin organization in situ to study genome organization, remodeling events, and architecture.

## Results

### Live cell Halo-Nanogold tagging using a bacterial toxin

To visualize endogenous proteins we developed a generalizable EM labeling strategy, Exogenously delivered via pore forming Streptolysin O (Slo) functionalized Nanoparticles (Exo-Slo-Nano). Once inside the cell, these functionalized nanogold particles become covalently conjugated to a target protein that is endogenously tagged, all within live cells. We use small gold nanoparticles (Au NP), 1.4 nm and 5 nm, with an attached Halo-Ligand to provide molecular specificity to the protein of interest, which carries an endogenous Halo-Tag. In the presence of the monovalent Nanogold-HaloLigand, the modified bacterial haloalkane dehalogenase, Halo-Tag, forms a covalent, irreversible linkage to its conjugate HaloLigand [32], yielding a covalently attached nanogold particle directly next to a target of interest (Fig 1a). HaloTag is relatively small (molecular weight 33 kDa) and has been used extensively as a genetically encoded molecular tag [32]. For visualization by live-cell fluorescence microscopy and CLEM methods, the 1.4 nm HaloLigand-Nanogold probes were also conjugated with Alexa fluorophores (Alexa Fluor 488 or 594 Alexa Fluor), yielding 1.4 nm-Halo-Alexa-Nanogold-488 (1.4 nm-HAN-488) or 1.4 nm-Halo-Alexa-Nanogold-594 (1.4 nm-HAN-594). Thus, these small probe designs possess (1) molecular specificity, (2) a high contrast nanogold moiety, and (3) are suitable tags for imaging from microns to nanometers (Fig. 1b). For these Au NP to be useful molecular tags in live cells, we sought to demonstrate that the probes can be robustly delivered into live-cells, detected in both fluorescence and electron microscopies, and recognize their targets of interest, and that the method is compatible with subtomogram analysis.

**Fig 1:**
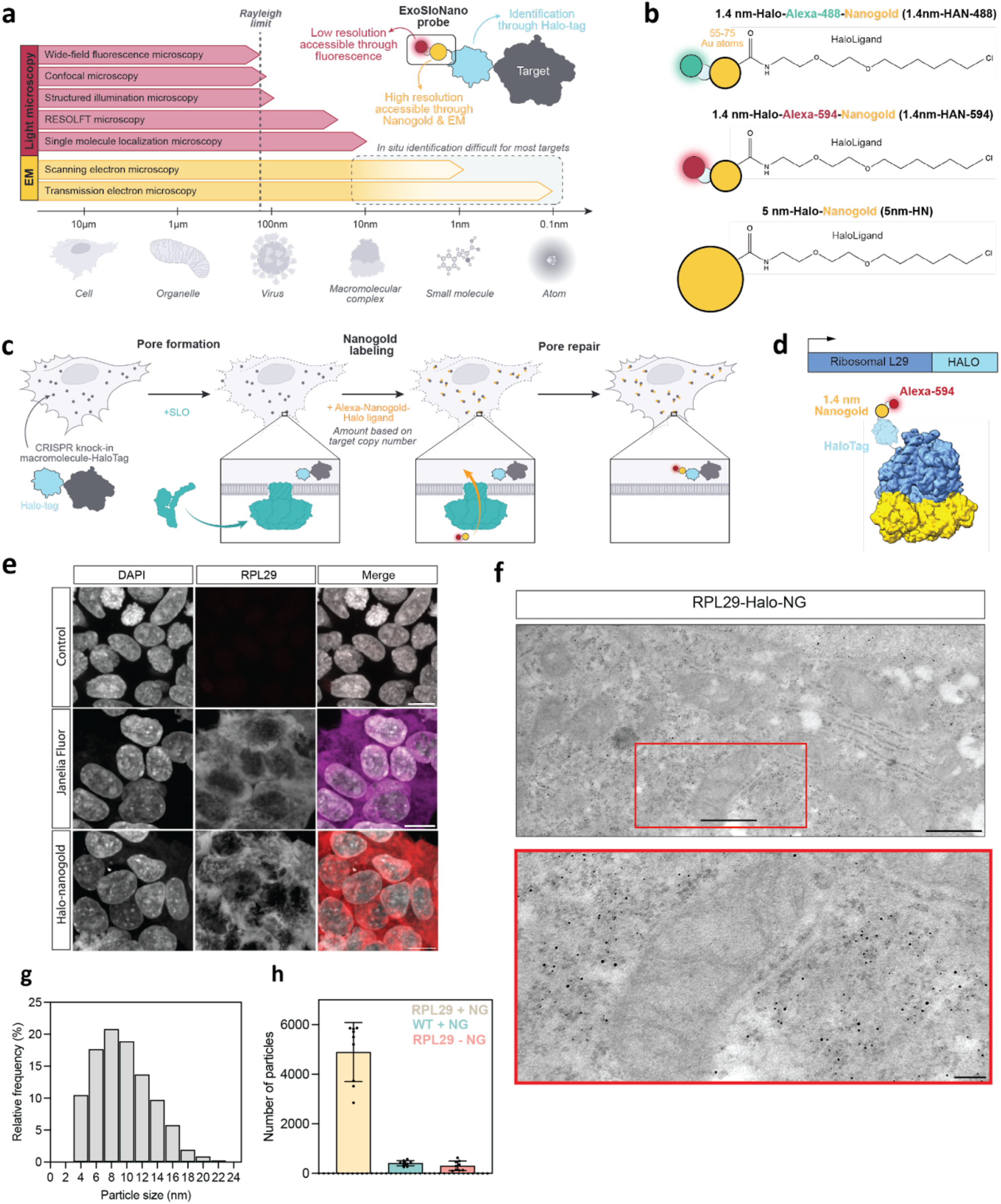
Nanogold probes and room-temperature TEM. (**a**) Need for more tools that span the meso and nanoscale. (**b**) Nanogold probes used in this study. 1.4 nm-Halo-Alexa-Nanogold-488 (1.4nm-HAN-488), 1.4 nm-Halo-Alexa-Nanogold-594 (1.4-nm-HAN-594), 5-nm-HaloNanogold (5-nm-HN). **(c)** Schematic of ExoSloNano: live cell delivery of exogenous nanoparticles by SLO. **(d)** Schematic of a covalent linkage of 1.4-nm-HAN-594 labeling of Ribosomal protein L29-Halo. **(e)** Live cell delivery of 1.4 nm-HAN-594 into RPL29-Halo cells: wildtype HEK 293 cells with nanogold delivery (top panel). RPL29-Halo cells treated with cell permeant Janelia Fluorophore (middle panel). Delivery of 1.4-nm HAN-594 into RPL29-Halo cells (bottom panel). **(f)** TEM micrograph of RPL29-Halo cells with gold enhancement of 1.4 nm HAN-594. Scale bar is 600 nm (top), 150 nm (bottom). **(g)** Histogram of the size distribution of gold particles after gold enhancement. N = 956 particles across two cells. **(h)** Plot of the number of nanogold particles found in the indicated conditions. N = 9 cells.

### Live cells recover from bacterial toxin treatment

Exogenous entry of nanogold particles into live cells was achieved by using the bacterial endotoxin Streptolysin-O (SLO) which forms small pores in the plasma membrane by binding to cholesterol [33]. Approximately 35-50 subunits oligomerize to form a pore ranging from 15-30 nm in diameter [34,35]. As a molecular tool, these bacterial pores have been used for exogenous delivery of membrane impermeant probes for super-resolution microscopy studies [36,37]. Following pore formation, the plasma membrane repairs after pore formation through an ESCRT-mediated mechanism [33,38]. Recovery post-SLO treatment is not cell type specific and has been shown in Chinese Hamster Ovary, NIH 3T3 fibroblast cells, Human Osteosarcoma (U-2 OS), Henrietta Lacks Ovarian (HeLa) cells, among others [36,39]. We monitored SLO-treated cells up to 100 hours post-treatment to confirm that cells recover (Extended Data Fig. 2b). We found that SLO treatment does not affect cell growth, and that growth is only mildly reduced by subsequent addition of nanogold (Extended Data Fig. 2a-c), with low levels of apoptosis that are comparable to untreated cells (Extended Data Fig. 2d). Finally, we found by room temperature EM that the cell’s ultrastructure is maintained after SLO treatment (Extended Data Fig. 2e). Together, these results indicate that SLO treatment provides broadly applicable method for nanogold delivery to live cells.

### Delivery of 1.4-nm-HAN-594 and enhancement with room temperature TEM

To test the ability of our probe to label specific proteins, we first targeted the ribosome, a large and abundant macromolecular machine that has been well characterized and beloved target for cellular cryo-ET studies [18,40,41]. To ensure that all ribosomes contained a HaloTag, we used a HEK293T knock-in cell line containing an endogenous C-terminal HaloTag7 on Ribosomal Protein L29 (RPL29) [42]. Previously, this cell line, RPL29-Halo was used as a marker to monitor ribosome turnover and degradation [42] as RPL29 is stably integrated, ensuring that each ribosome carries a HaloTag. We used this cell line to validate that our nanogold probes label the intended target using room-temperature TEM and cryo-ET.

We first demonstrated nanogold labeling using resin-embedded room temperature TEM. To this end, we delivered Halo-Alexa 594-1.4-nm Nanogold (1.4 nm-HAN-594) to HEK 293T L29-HaloTag cells (RPL29-Halo) (Fig. 1d). Wildtype cells treated with SLO and 1.4 nm-HAN-594 do not show fluorescence (Fig 1e, top panel). In contrast,RLP29-Halo cells treated with SLO and 1.4 nm-HAN-594 show fluorescence within the cytosol (Fig 1e, bottom panel) that is comparable in intensity and spatial distribution to the fluorescence of a commonly used cell permanent Janelia fluorophore which contains a HaloLigand (Fig 1e, middle panel). This shows that nanogold labeling is specific to the presence of the Halo Tag, and suggests that live nanogold tagging may not perturb protein localization. Following nanogold delivery, cells were fixed, exposed to silver which binds to and enhances 1.4 nm-HAN-594, which subsequently increases in diameter. Then, cells were resin-embedded, stained with a low percentage of osmium tetroxide for contrast (0.7%), and sectioned (Fig. 1f). Visualized by room temperature EM, the enlarged 1.4-nm-HAN-488 have a mean diameter of 9.6 nm, and show a broad, uniform distribution of nangold throughout the cytoplasm and endoplasmic reticulum (Fig 1f). Nanogold enhancement was consistent with the presence of RPL29-Halo, yielding robust cellular recruitment in the presence of the Halo-tagged target; 5e4 particles were detected in a 10.1 um^*3*^ volume (Fig 1g, Extended Data Fig. 3a). When 1.4-nm-HAN-594 was supplied in cells without a HaloTagged protein (wild type cell, WT), nanogold enhancement and detection was negligible (Fig 1h). Similarly, nanogold enhancement and detection was minimal in RLP29-Halo cells when 1.4 nm-HAN-594 was not supplied (Fig. 1h, Extended Data Fig. 3a). Using an established protocol to determine cellular copy number [43], we determined that there are approximately 1.7e6 ribosomes per HEK RPL29-Halo cell (Extended Data Fig. 3e). Extrapolating the number of enhanced nanogold particles detected from a 10.1 um^*3*^ volume to the entire volume of a HEK L29-Halo (5282.5 um^*3*^) cell yields 2.6e6 ribosomes per cell (Extended Data Fig 3b-d). Overall, this is consistent with the saturation of L29-Halo binding sites due to the delivery and enhancement of the 1.4 nm-HAN-594 probes (see below).

### Detection of 5-nm Halo-Nanogold by cryo-ET

We next sought to visualize individual ribosomes with bound nanogold by cryo-ET. The 5 nm-Halo-Nanogold (5nm-HN) are readily visible in cryo-ET data, and were utilized to optimize parameters for data collection, tilt series alignment, and subtomogram analysis of in situ tomograms by cryo-ET. 5-nm-HN was delivered to live RPL29-Halo cells as described above. Cells recovered overnight, then were seeded onto EM grids, vitrified by plunge freezing, FIB, milled, and tilt series were collected (Extended Data Table 1). Individual 5-nm-HN could be readily identified directly adjacent to ribosomes (Fig 2c, 2d). Automated particle picking of 13,748 ribosomes across 46 tomograms and subsequent subtomogram alignment and averaging yielded a bright mass (Fig 2c) with a size consistent with a 5-nm nanogold particle. Unsurprisingly, this indicated that the high-intensity region in the subtomogram corresponding to the high-mass density of the nanogold particle drives the alignment.

**Figure 2:**
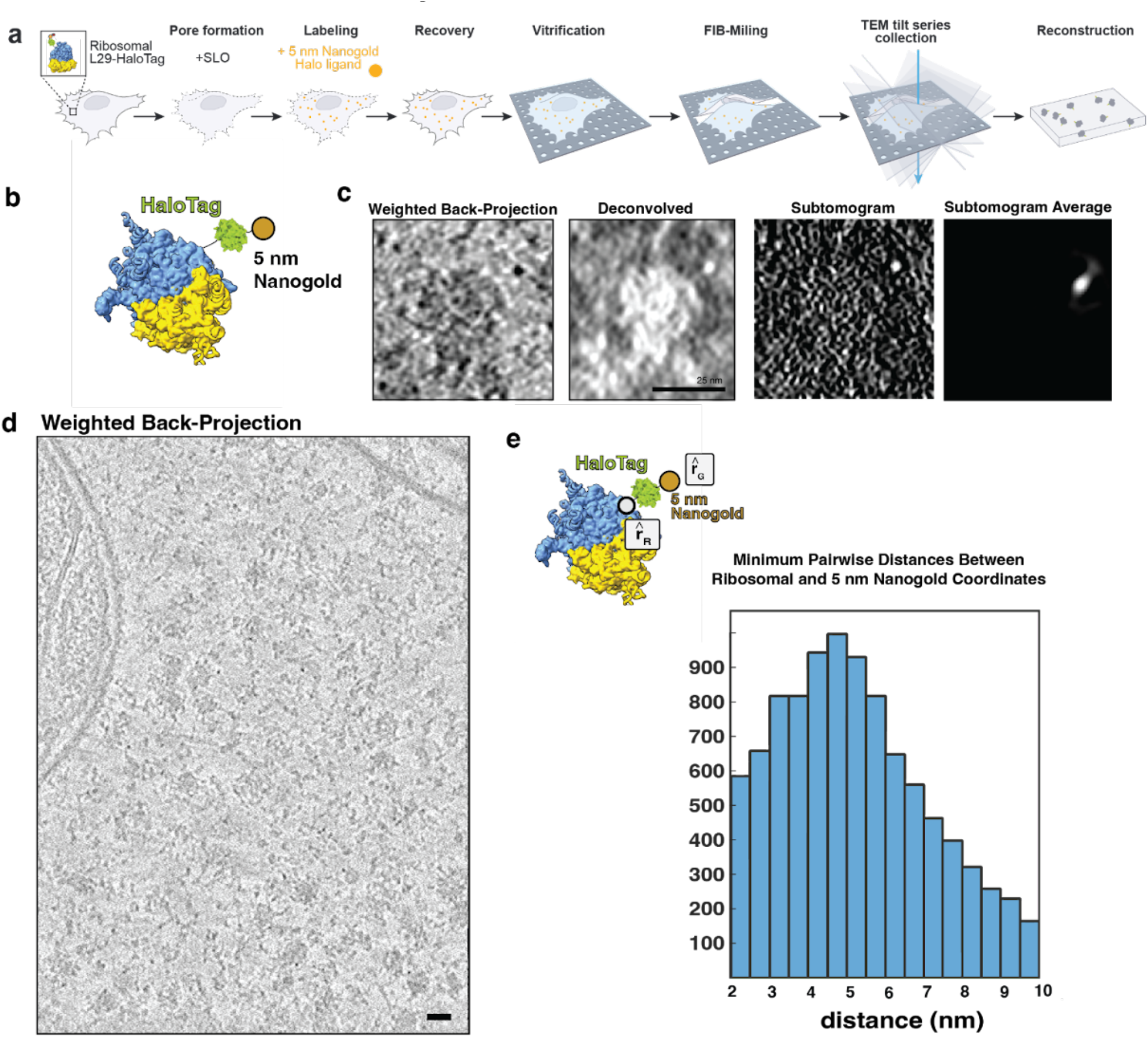
Cellular cryo-ET of 5 nm-HN labeled ribosome. **(a)** Schematic of live cell delivery of 5 nm-Halo-Nanogold (5nm-HN), vitrification, cryo-FIB-milling, cryo-ET, and tomographic reconstruction to generate 3-D volumes. (**b**) Schematic of a 5 nm-NH labeled ribosome, white dot represents the C-terminus of L29 that defines r_R_, and the center of the nanogold defines r_G_. (**c**) Central slice (8 Å/pixel) of an individual ribosome labeled with 5 nm-NH by weighted back-projection (contrast is as in original data, where dark pixels correspond to high occupancy, i.e., black-on-white), the same subtomogram deconvolved, its corresponding normalized subtomogram, and the subtomogram average from 13,748 ribosomal picks, dominated by alignment of nanogold intensity (contrast inverted, i.e., white-on-black). Scale bar is 25 nm. (**d**) Central slice of a weighted-back projection of the whole tomogram, averaged over 10 slices, scale bar is 20 nm. **(e)** Schematic ribosomal distance of ribosome and nanogold. Pairwise distances between (r_R_) and 5 nm-HN (r_G_).

To determine labeling efficiencies, the pairwise distances between these 13,748 particles were determined between nanogold coordinates and the center of the refined ribosome coordinates. The nanogold coordinates were obtained based on the initial alignment including the nanogold signal, whereas the ribosome coordinates were obtained from the latter alignment without it. The distribution of distances yielded a distribution in which the average distance is 16.8 nm (+/-3 nm) between the center of mass of the ribosome and the center of mass of the 5 nm nanogold. Then, 12.9 nm, the distance from the center of a ribosome to the last ordered residue of L29, was subtracted, and those values were plotted. Of the 13,748 picks, 9,567 particles, or approximately 69.5% percent, were determined to be within a distance of 5.22 nm (+/-1.9 nm) (Fig. 2e). These distances agree with the manual identification of distances between individual ribosome and nanogold pairs (Extended Data Fig. 4).

### Detection of 1.4 nm-HAN-488 by cryo-ET

In order to test a smaller probe, optimization of data collection conditions and preprocessing steps using the 5 nm-HN were applied to directly visualizing ribosomes labeled with 1.4-nm-HAN-488 by cryo-ET. Briefly, 1.4 nm-HAN-488 was delivered to RPL29-Halo cells via SLO, 18 hours later, cells were seeded onto EM grids, samples were then vitrified after adhesion 4-6 hours, FIB-milled and tilt series were collected (Fig. 3a and Extended Data Table 1). Data was collected at sufficient magnification (1.068 Å/pixel) to ensure that there would be enough pixels to represent an individual nanogold moiety (14 Å, ∼13 pixels). As with 5 nm-HN, 1.4 nm-HAN-488 could be visualized directly adjacent to the ribosome within a tomogram (Fig. 3e). The average pairwise distance from RPL29 and the nanogold was 5.1 nm (+/-2.4 nm) (Fig. 3g). Notably, these data show that nanogold particles of multiple sizes can be employed in in situ cryo-ET experiments in order to facilitate multiplexing, i.e., the tagging of multiple cellular components simultaneously. To demonstrate that these particles can be distinguished and thereby enable multiplexing, we deposited nanogold particles on EM grids and quantified their diameter. This showed clearly separated size distributions with a mean diameter of 1.52 nm (+/-0.359 nm) and 5.62 nm (+/-0.911 nm) for 1.4 nm and 5 nm-HN, respectively (Extended Data Fig. 1a-d).

**Figure 3.**
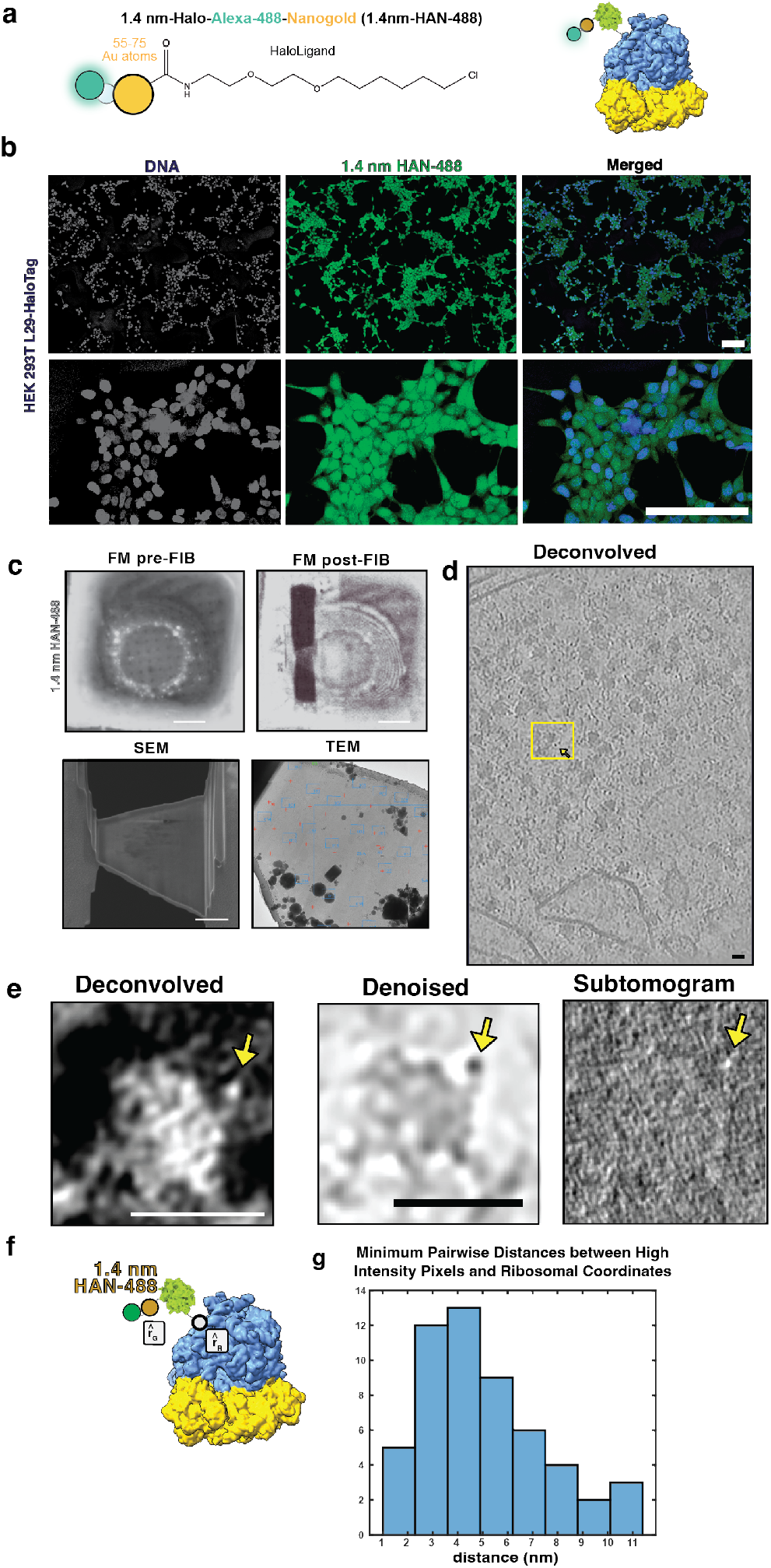
Ribosomal labeling with 1.4 nm-HAN-488. **(a)** Schematic of probe used to label the ribosome **(b)** HEK293T RPL29Halo SLO-treated cells with 1.4 nm-HAN-488. Top: scale bar is 100 microns. Bottom: scale bar is 50 microns. **(c)** Fluorescence microscopy (FM) of a grid pre-FIB and post-FIB milling, SEM-view of the lamella, and TEM view of the lamella. **(d)** Central slice of a deconvolved tomogram (contrast is black-on-white), scale bar is 20 nm. **(e)** Central-slice of an individual ribosome labeled with 1.4 nm-HAN-488 shown as a deconvolved, denoised, and a normalized subtomogram (4 Å/pixel). Scale bar is 25 nm. **(f)** Schematic of 1.4nm-HAN-488 labeled ribosome to represent the distance from ribosomal subunit L29 (r_R_) and 1.4nm-HAN-488 (r_G_). **(g)** Pairwise distances between ribosome coordinates and 1.4 nm nanogold coordinates.

### Expanding the proteome to challenging targets

We developed this method generally to expand the proteome available to cellular EM. However, our ultimate goal was to tackle targets that have proven challenging to available methods. Notably, the nucleus has largely remained an uncharted territory for *in situ* structural cell biology. Even for ultrastructure studies using roomtemperature EM, the nucleus has been referred to as “the dark side of the cell”, largely due to the challenge with antibodies penetrating into nucleus [44]. Great progress has been made in imaging chromatin by room temperature EM [9] and cryo-ET [45], but it has so far not been possible to discern histone variants and nucleosome subpopulations, i.e. we currently cannot distinguish heterochromatin vs euchromatin beyond inferring it from the degree of compaction. To our knowledge, no exogenous, e.g., antibodies or endogenous, e.g., small genetically encoded proteins visible in cryo-EM, have been able to specifically target and label histone-specific nucleosomes with minimal disruptions to the chromatin landscape.

The role of chromatin architecture in genome function and maintenance is a very active field of research. Powerful, but indirect methods such as chromosome conformation capture assays, including Hi-C [46], map DNA interactions at a range of scales (0.1-1 Mb), either from an individual cell to a population of cells [47]. A major limitation is that chromatin capture experiments only capture genomic DNA interactions, i.e. nuclear macromolecular complexes are lost. To complement chromatin capture experiments, in situ structural biology has the potential to map spatial relationships within native chromatin landscapes.

Genome organization relies on the incorporation of canonical, non-canonical nucleosomes and their associated epigenetic markers, among other factors [48]. Canonical nucleosomes are octomers containing two copies of replication-coupled histone proteins H2A, H2B, H3.1, and H4, wrapped by ∼145-147 bp of DNA [49]. Variants of H2A, H2B, and H3 allow greater complexity of epigenomic regulation, as their incorporation is specific to cell type, developmental and functional state, among other aspects [50]. In mammals, there are 19 variants of H2A alone [51]. Histone variants play important roles in genome maintenancei.e. the phosphorylation of H2A.X labels sites of dsDNA breaks for DNA repair processes [52], H3 variant CENP-A localizes to centromeric DNA [53], and H2A.Z is incorporated around transcription start sites [54,55]. Nucleosomes from different species and nucleosomes containing histone variants have high structural homology [56], with <1 Å^*2*^ root-mean-square deviation when the backbone of the structures are compared. The largest variation in their structure is due to the “breathing” of the linker DNA near to the entry and exit of the nucleosome [57]. Thus, even with the advent and progress of template matching methods for cryo-ET data, it will be difficult to discern nucleosome subpopulations within the nucleus. As a test for our method, we chose macroH2A (gene name H2AFY), a histone with broad spatial distribution across the nucleus (Fig. 4b), but enriched at heterochromatin regions [58], lamin-associated domains (LADs) [59], and most notably, the inactive X-chromosome [60].

**Figure 4:**
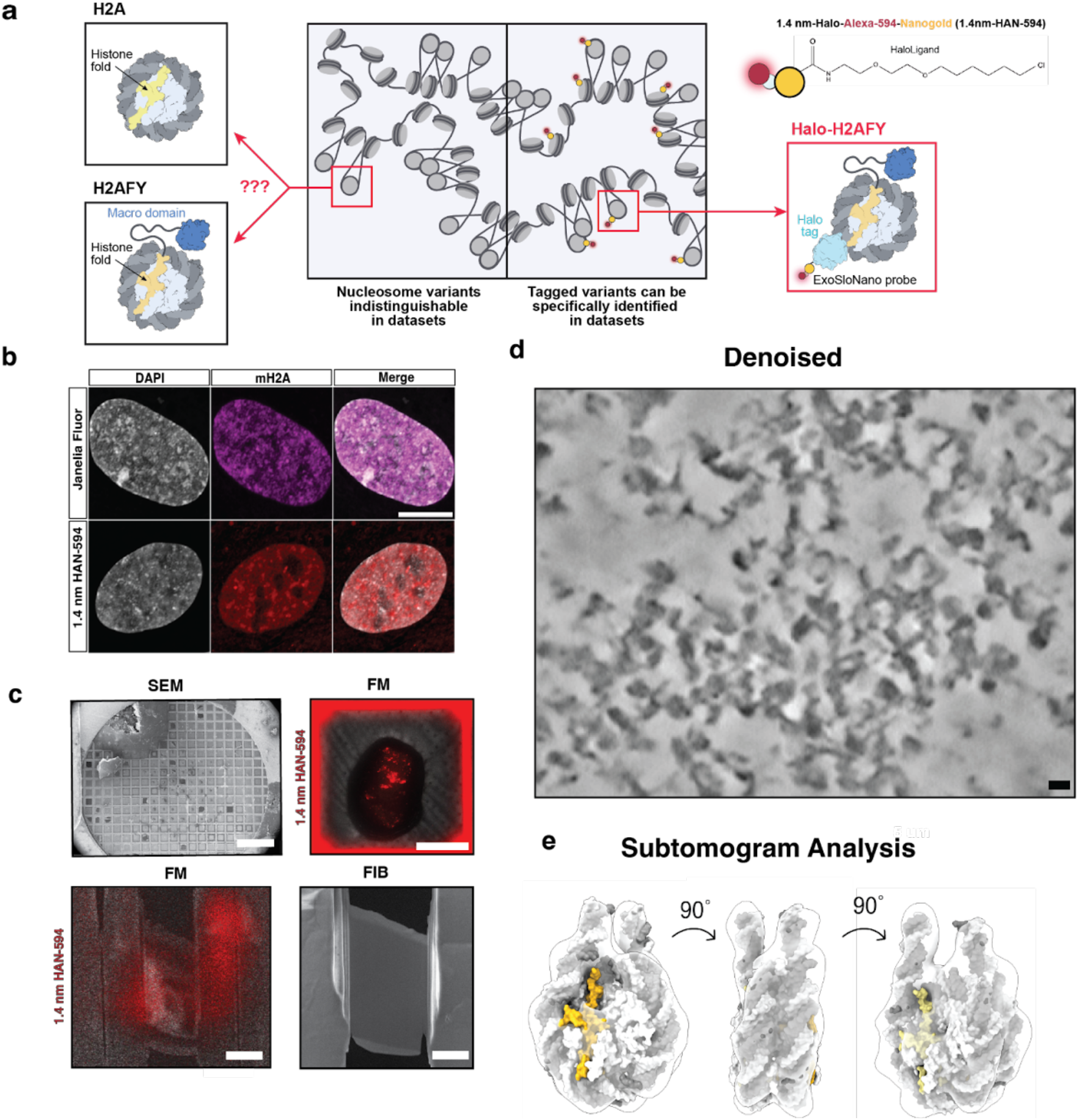
Nuclear delivery of 1.4 nm-HAN-594 to target heterochromatin. **(a)** Protein domain schematic of relevant histone variants **(b)** Top panel: RPE1 Halo-mH2A cells labeled with membrane permeant Janelia Fluor which freely diffuses into cells and recognizes the HaloTag. Bottom panel: RPE1 Halo-mH2A cells labeled with 1.4-nm-HAN-594 delivered via SLO **(c)** Cryo-FIB-ET workflow: RPE1 Halo-mH2A cells were grown on an EM grid (scale bar 100 um), cryo-fluorescence image before FIB-milling (scale bar 30 microns), during milling (scale bar 10 microns), and post-milling (scale bar 5 microns). **(d)** Denoised and deconvolved in situ tomogram of chromatin, scale bar 10 nm. **(e)** Subtomogram analysis of all nucleosomes.

Histone variant H2AFY encodes macroH2A1 (mH2A) and its splice variants, macroH2A1.1 and macroH2A1.2 [61]. MacroH2A variants contain a histone fold, a lysine-rich linker domain, and a C-terminal macrodomain [62]. Although the histone fold of mH2A has 64% sequence identity to canonical H2A [63], nucleosomes containing single or double copy of the mH2A histone have high structural similarity to the canonical nucleosome [64]. The C-terminal macrodomain is a globular domain that binds ADP-ribose and polymers of ADP-ribose. Notably, macrodomains are present in all domains of life-archaea, bacteria, eukaryotes, and viruses [65].

MacroH2A plays an important role in maintaining heterochromatin architecture [66]. It has been found at transcriptionally silenced genes, found to co-localize with heterochromatin marker H3K27me3 [66], and Polycomb Repressive complex 2 [67], among other markers of silenced chromatin. The presence of mH2A is an epigenetic barrier to somatic cell reprogramming [66,68–70]. Finally, mH2A and its isoforms are tumor suppressors-the loss of mH2A correlates in more aggressive cancer phenotypes [71–74] and low expression of macroH2A1.1 is associated with poor prognosis in multiple cancers (colorectal, lung, astrocytoma, and breast cancers), [72,75–77]. All of this points to the importance of mH2A in heterochromatin maintenance.

Although mH2A plays an important role in chromatin architecture and maintenance, we currently do not have a way to identify and spatially resolve mH2A-containing nucleosomes within the nucleus. We reasoned that 1.4nm-HAN-594 and cryo-ET would be ideal for probing histone variants, as the small probe elicits minimal perturbations (Extended Data Fig. 7). To obtain endogenous levels of Halo-tagged macroH2A, cultured human Retinal Pigment Epithelial (RPE1), a primary female derived cell line, was CRISPR engineered to bear an N-terminal HaloTag and a V5 Tag epitope at the endogenous H2AFY locus (Fig. 4a, Extended Data Fig. 5). The endogenous integration of HaloTag was verified by sequencing (Extended Data Fig. 5).

### Quantification of an endogenous target and nanogold delivery

In order to match the internal cellular concentration of the target and mitigate unlabeled targets and unbound nanogold, appropriate exogenous nanogold concentrations must be delivered to the cell. This requires knowledge of the target’s cellular copy number and quantification of probe delivery. We used the earlier mentioned established protocol to determine the copy number of mH2A in RPE1 cells [43].

From this, we determined that there are approximately 7e5 copies (+/-2.3e5) of mH2A within the nucleus (Extended Data Fig 6). To quantify nanogold delivery into the cell, RPE1 mH2A-Halo cells were exposed to SLO and 1.4 nm-HAN-594, and the fluorescence intensity per cell was measured against a dilution series of the 1.4 nm-HAN-59 probe. The internalized concentration of 1.4 nm-HAN-594 was determined to be 6.6e5 copies (+/-1.05e5) per cell (Extended Data Fig. 6).

### Nanogold labeling does not perturb chromatin architecture

Cultured RPE1 Halo-mH2A cells were exposed to SLO for live-cell labeling with 1.4 nm-HAN-594. The fluorescence of 1.4 nm-HAN-594 shows nuclear localization, comparable to a control cell labeled with a cell permeant fluorophore containing a HaloLigand (Janelia Fluor). In both samples, there is an enrichment in the nucleus and the chromatin dense regions, consistent with the presence of mH2A in compact chromatin (Fig. 4b). To confirm that 1.4 nm-HAN-594 tagging does not perturb chromatin ultrastructure, chromatin organization was visualized by an established protocol for studying chromatin by room temperature TEM, Chromatin Electron Tomography (ChromEMT). Briefly, fixed RPE1 Halo-mH2A cells, in the presence or absence of 1.4-nm-HAN-594, were treated with the DNA dye DRAQ5. This was photo-oxidized using 630 nm excitation and diaminobenzidine (DAB), leading to the formation of AB polymers on the chromatin surface which are visualized by EM via binding of Osmium Tetroxide [9]. Dual axis tomography datasets were collected and chromatin diameter and packing density was quantified (Extended Data Fig. 7), showing similar distributions between conditions, suggesting that nanogold tagging does not perturb global chromatin packing or ultrastructure.

### Subtomogram analysis of mH2A labeled 1.4 nm-HAN-594

As there are two copies of H2A within the nucleosome octamer, a single copy could be replaced with an mH2A variant (creating a hybrid or heterotypic nucleosome), or both copies of H2A could be replaced (Fig. 5a). Single and double incorporations of mH2A are highly structurally homologous to a canonical nucleosome. Intriguingly, it was found that there is a rearrangement of the L1-L1 interface in mH2A containing nucleosomes [78]. The L1-L1 interface is the only region where the two H2A-H2B dimers interact [49]. In vitro conditions, this results in an increased thermodynamic stability of hybrid nucleosomes [79]. As far as we are aware, how mH2A is incorporated into nucleosomes inside the cell has not yet been determined.

**Figure 5:**
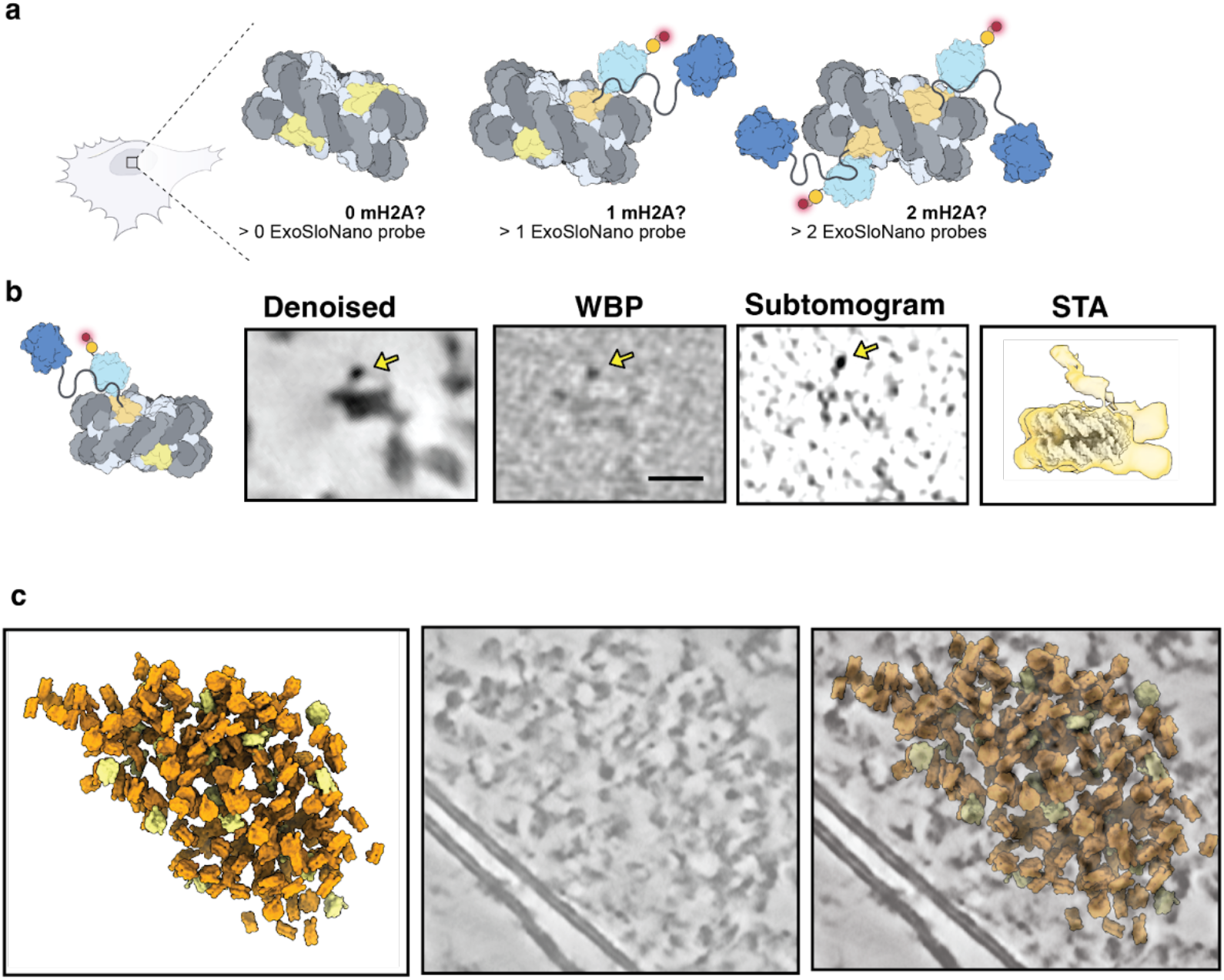
Isolating macroH2A-nucleosomes from in situ cryo-ET data. **(a)** Diagram of the possible macroH2A incorporations per nucleosome and the number of ExoSloNano probes **(b)** Single nucleosome from of RPE1 mH2A-Halo cell treated with 1.4-nm-HAN-594 of the central slice at 8 A/pixel from Denoised, Weighted Back Project (WBP), a normalized subtomogram, and from subtomogram analysis (STA). Scale bar is 10 nm. Yellow arrows point to a density consistent with a 1.4 nm nanogold moiety. Subtomogram analysis on nanogold-labeled nucleosomes. **(c)** Sub-tomograms mapped back to the original tomogram containing labeled nucleosomes (yellow) and unlabeled nucleosomes (orange).

In order to isolate nanogold-labeled nucleosomes, all nucleosomes were picked by template matching on deconvolved tomograms, and subtomogram averaging was performed, resulting in an 18 Å reconstruction. Then, 3D classification with local angular sampling using an inverse mask around the nucleosome was performed. Particles were selected for those containing a globular density consistent with the size of 1.4 nm nanogold probe (Fig. 5b). The average contains a diffuse density corresponding to a flexible nanogold probe 5.5 nanometers above a nucleosome. Notably, either zero or one nanogold is present in the nucleosome averages, in agreement with the in vitro data suggesting that in nucleosomes containing mH2A, hybrid mH2A composition is the most stable, i.e., we did not observe nucleosomes with two mH2A histones. Overall, this demonstrates that macromolecules of limited copy number can be recovered using our ExoSloNano approach.

## Discussion

We present a method for delivering, labeling, and visualizing macromolecules in their native context at high spatial resolutions by two different modalities. Fluorescently-labeled nanogold can be used to label macromolecules in live cells to be visualized by fluorescence microscopy and be further identified in EM studies. This strategy is compatible with live cells and does not alter ribosomal localization or heterochromatin architecture. Importantly, this only requires the addition of a HaloTag to the protein of interest; the subsequent covalent linkage to a nanogold particle occurs within the cell. Notably, this tagging approach does not require complex molecular assemblies or sensitive in vivo reactions. Finally, it can be visualized in in-cell cryo-EM, here shown for in situ cryo-ET. Besides the power of protein localization within intact cellular networks, the presence of nanogold moieties should act as internal fiducials, improving cryo-ET data reconstruction steps such as motion correction and tilt series alignment, and thereby potentially facilitate template matching of smaller targets inside the cell both in 2-D and 3-D.

We show how 1.4 nm and 5 nm nanogold probes can be detected by cryo-ET. We demonstrate the feasibility of our approach for future multiplexed labeling, wherein gold particles of different sizes are conjugated to self-ligands, here shown, but not limited to HaloTag. This method can readily be expanded to use SNAP tags, click chemistry, labels that recognize epigenetic markers, or any other biological or chemical moiety that can be adapted for labeling; adaptation to nanogold labeling and delivery should be easily adaptable to further extend the toolkit. The ExoSloNano approach presented here will allow for detailed interrogation of complex interactions within the cell and the nucleus, and likely other organelles. Ideal implementation would utilize cell lines with an endogenously-tagged macromolecule of interest of which there is a prior estimation of copy number. The methods presented here would complement the progress within the chromatin organization field and provide more tools for probing chromatin organization, in vitro and in situ, facilitating advancement in studying genome organization, and genome remodeling events. Combined with recent advances in resolving and analyzing chromatin by cryo-ET [41,45,80,81], our labeling strategy has potential to directly visualize chromatin regulators as they interface with chromatin in situ.

Currently, in situ cryo-EM is not even close to the theoretical limitations on detection of molecules of sub-100 kDa [82,83]. To truly expand what we can answer by cellular cryo-ET, we need more tools that are orthogonal in their labeling methodologies. This method in conjunction with existing probes such as DNA Origami Signpost, Ferritag, and GEM will further empower cell and structural biologists to identify macromolecules across scales [12–14]. We encourage academic and commercial chemistry efforts to lead the development of homogenous and monovalent nanogold particles of additional sizes. Eventually, swapping nanogold with a STEM-EELS compatible probe will further expand in situ molecular identification of novel targets inside cells by cryo-ET, which will enable multiplexing of targets using multiple size tags and EELS signatures. We hope that the method presented here empowers the cellular cryo-ET to expand the accessible proteome to advance the potential of structural cell biology.

## Acknowledgements

We thank Paul Selvin for insightful initial conversations about using SLO for probe delivery, Niko Grigorieff, Bing Ren, Tom Laughlin, and members of the Villa lab for helpful discussions. We thank the following imaging facilities at UCSD: Neuroscience Imaging Center, the Nikon Imaging Center at UC San Diego, and the UCSD Cryo-Electron Microscopy Facility, which was built and equipped with funds from UCSD and an initial gift from the Agouron Institute. We thank the UCSD Physics Computing for computational support. M.K.R. and E.V. are Howard Hughes Medical Institute Investigators. This work was supported by an NIH NCI K00 grant (CA223029 to L.N.Y.), an EMBO long-term postdoctoral fellowship ALTF #902-2019 (to A.S), an NIH K99 HD112607-02 (to A.S), an NIH DP2 (DP2-GM-123494 to E.V.), a Pew Scholar Award (to E.V.), and an NSF DBI 1920374 (to E.V.).

Molecular graphics and analyses were performed in part with UCSF ChimeraX, developed by the Resource for Biocomputing, Visualization, and Informatics at the University of California, San Francisco, with support from National Institutes of Health R01-GM129325 and the Office of Cyber Infrastructure and Computational Biology, National Institute of Allergy and Infectious Diseases. Computational support from the PCF. HEK 293T L29-Halo was a kind gift from Heesson An. This paper was typeset with the bioRxiv word template by @Chrelli: www.github.com/chrelli/bioRxivword-template

## Author contributions

L.N.Y., A.S. and E.V. conceived the research. L.N.Y. and A.S. designed the experiments. A.S. generated the RPE1 H2AFY cell line. L.N.Y. and F.S. prepared the samples for cryo-FIB-ET. L.N.Y. and F.S. prepared cryo-lamellae. L.N.Y. collected the cryo-electron tomography data. J.H. established PACE-tomo at UCSD. M.R. created the schematics. L.N.Y carried out tomographic reconstruction, subtomogram averaging, and image analysis. H.Z. analyzed the chromatin data. A.S. prepared, imaged, and analyzed resin embedded samples. L.N.Y., A.S. and E.V. wrote the manuscript with contributions from all authors.

## Materials and Methods

### Cell culture

Retinal Pigment Epithelial (RPE1) H2AFY and WT RPE1 cells were cultured in DMEM with 4.5 g/L D-Glucose, and L-Glutamine (Gibco, 11965-092) supplemented with 10% Fetal Bovine Serum (Gibco One Shot, A3160401) and Antibiotic-Antimycotic (100 U ml^−1^ penicillin, 100 µg ml^−1^ streptomycin, Gibco, 15240062). U2OS C32 Halo-CTCF cells were cultured in DMEM (low glucose, GlutaMAX Supplement, pyruvate (Fisher Scientific, catalog number: 10567014) supplemented with Antibiotic Antimycotic (Gibco, 15240062) and 10% Fetal Bovine Serum (Gibco One Shot, A3160401). HEK 293T L29-HaloTag cells were cultured in DMEM with 4.5 g/L D-Glucose, and L-Glutamine (Gibco, 11965-092) supplemented with 10% Fetal Bovine Serum (Gibco One Shot, A3160401) and Antibiotic-Antimycotic (100 U ml^−1^ penicillin, 100 µg ml^−1^ streptomycin, Gibco, 15240062). Cells were cultured at 37°^C^ and 5% CO_2_. All cell lines were tested regularly for mycoplasma by PCR (Myco-Sniff mycoplasma PCR detection kit).

### Validation of CRISPR knock-in HEK RPL29-Halo

CRISPR knock-in HEK L29-Halo cells were validated through Sanger sequencing of the endogenous RPL29 gene. The linker connecting RPL29 and HaloTag7 is GSGGSAEIGTGFPFDPHYVEVLGER. The gene product was verified by a Western blot against HaloTag, and fluorescence microscopy.

### HaloTag CRISPR-Cas9 mediated endogenous genome editing

To generate the targeting plasmid, gibson assembly was used to clone a V5 epitope, followed immediately by a HaloTag into pUC19. The V5-Halo tag is upstream of H2AFY and is joined using a GDGAGLIN-linker. To generate the gRNA plasmid, the gRNA sequence (5’ CACCGCCCAC-CGCGGCTCGACATGG 3’) was cloned into PX458 using Bbs1 digestion and established protocols [84]. To generate an endogenous H2AFY-Halo-Tag cell line, 1 million hTERT immortalized Retinal Pigment Epithelial (RPE1) cells (ATCC CRL-4000TM) were electroporated with 7.5 ug of tar-geting plasmid and 6ug of PX458 using the Neon transfection system (Invitrogen). Following electroporation, 50nM JF-Halo ligand (647, Promega) was added to cells and fluorescence activated cell sorting (FACs) was used to purify cells with a Halo-tag knockin. Single cell colonies were genotyped using the following primers, which were designed to amplify the N terminal region of H2AFY. PCR fragments were separated by gel electrophoresis, obtaining the expected band shift, and bands were gel purified and sequenced using Sanger sequencing to confirm V5-Halo knockin.

Fwd primer: 5’ TCCAGCGGAGTGCATCACC 3’

Rev primer: 5’ TTCTTGATGTACCGCAGCATCCG 3’

### Flow Cytometry and Protein Abundance

The absolute protein abundance of H2AFY was determined through flow cytometry when compared to a known standard, U2OS C32 Halo-CTCF cells were cultured in DMEM supplemented with 1 g/L Glucose and 110 mg/L sodium pyruvate (Gibco, 10567-014) [43]. U2OS C32 HaloTag-CTCF cells were a gift from Robert Tijan’s lab (UC Berkeley). U2OS C32 Halo-CTCF cells and RPE1 H2AFY-Halo cells were cultured in 6 well plates to 90% confluency, to which 1 uM HaloTag TMR ligand (Promega, G825A) was supplied for 30 minutes. Cells were washed twice with 1x DPBS, trypsinized with 0.5% Trypsin (Gibco 15400054), gently resuspended in HBSS (Gibco 14025134), spun down and gently resuspended in HBSS with 50 ug/ml deoxyribonuclease I from bovine pancreas (Sigma D4263). For each sample, >10,000 events/rates were analyzed on a BD Biosciences LSR-II flow cytometer using a 561 nm excitation laser line and a 582 / 15 band pass filter cube. Absolute protein abundance was determined using an established protocol [43].

### Streptolysin O (SLO) treatment

HEK 293T WT and RPL29-Halo cells were cultured on fish gelatin coated 24-well plates, when cells reached 70%-80% confluency, they were treated with 40-100 units of streptolysin O (SLO) from Streptococcus pyogenes (25,000–50,000 U, Sigma-Aldrich, S5265). Prior to use, SLO was activated with 10 mM TCEP pH 7.0 for 20 minutes at 37 °C. Cells were exposed to SLO for five minutes, then gently washed two times with 1x DPBS, then exposed to CF® Dye 488 Dextran Anionic and Fixable (Biotium 80110), 10kDa, 40 kDa, or 250 kDa. Cells were incubated in recovery media consisting of OptiMEM, 10% FBS, 1 mM CalCl_2_, 1 mM ATP, 1 mM GTP, 2 mM Glucose. In experimental conditions following SLO treatment optimization, cells were cultured to 70%-80% confluency and treated with 40-100 units of SLO (Sigma-Aldrich, S5265) for five minutes, then gently washed two times with 1x DPBS, cells were exposed to 1.4 nm HAN-488, 1.4 nm HAN-594, or 5 nm-HN for eight minutes on a rotating platform, then gently washed two times with 1x DPBS and left to recover in recovery media consisting of OptiMEM, 10% FBS, 1 mM CalCl_2_, 1 mM ATP, 1 mM GTP, 2 mM Glucose.

### Live cell imaging post-streptolysin O treatment

Cells were imaged for 60 hours following SLO treatment to monitor cell survival. Cells were incubated in phenol-free OptiMEM media supplemented with 10% Fetal Bovine Serum, 1% Antibiotic Antimycotic (Gibco), 1 mM CaCl_2_, addition of Propidium iodine (0.5 ug/ml, BD Biosciences), and fluorescently labeled Annexin V-AF647 (Biolegend) was supplied to monitor cell death and apoptosis, respectively. Images were acquired on a Nikon Eclipse Ti2-E equipped with a Qi-2 camera and using Nikon Elements 5.02.02 software. Cell survival and cell growth was imaged over 60 hours at 37°C with 5% CO_2_ in stage top incubator (Okolab H301 Bold Line). Three non overlapping 3.125 mm^2^ zones within each well were imaged by DIC, 554/609, and 618/698 nm using a SpectraX light engine (Lumencor) with individual LFOV filter cubes (Semrock). Images were acquired every 20 minutes. Cell viability was determined by the ratio of live to dead cells.

### Nanogold-HaloLigand probes

1.4-nm Halo-Alexa-Nanogold-594, 1.4-nm Halo-Alexa-Nanogold-488, and 5-nm Halo-Nanogold conjugates were custom ordered from Nanoprobes, Inc. 1.4 nm HAN-488 and 1.4 nm HAN-594 arrived as 30 nmol, lyophilized from 1 mL of 0.02 M sodium phosphate buffer, pH 7.4, with 0.15 M sodium chloride, and was rehydrated in 1 mL of ultrapure water to make 30 uM stocks, 50 ul aliquots were stored at -80 C in PCR tubes.

### Quantification of Fluorescent Nanogold delivery

A dilution series of 1.4 nm HAN-594 was generated, the fluorescence intensity was measured (Molecular Devices SpectraMax i3x). The fluorescence intensity versus the number of 1.4 nm-HAN-594 molecules was plotted and a linear equation was fit. The fluorescence intensity was measured from SLO-treated and 1.4 nm-HAN-594-delivered cells. Background signal from SLO treated cells, but not treated with 1.4 nm HAN-594, was subtracted. Using the linear fit, the number of nanogold molecules that were internalized was determined which was divided by the total number of cells to determine the number of 1.4 nm HAN-594 molecules that were delivered to a RPE1 Halo-mH2A cell (Extended Data Fig. 6b). This value was determined by a bulk measurement, serving an approximation of the number of 1.4 nm-HAN-594 molecules delivered to each cell.

### InCuyte imaging

Following streptolysin O (SLO) treatment and nanogold labeling cells recovered for 4 h in recovery media, which was then replaced with normal media. Cells were placed in an inCyute microscope (Satorius), maintained at 37 degrees and 5% CO2. Four images were acquired using a 10x objective in each well (3 replicates per condition) every 3 h for a period of 4 days. Growth curves were calculated by cell confluence and were pooled for each between wells and replicates.

### Culturing cells on Micropatterned EM grids

Electron microscopy quantifoil grids (R 1/4 Au 200-mesh) were plasma cleaned on a Pelco easiGlow plasma cleaner for 1 minute at 20 mA on both sides. Ten ul of PLL-PEG (0.5ug/ml) in 10 mM HEPES buffer, pH 7.4 was applied for 45 1 hour at room temperature to an EM grid, then washed four times with 1x PBS and then 3 ul of the photoactivatable reagent PLPP (Avéole) was applied to each grid just prior to patterning. EM grids were imaged on a Eclipse Ti2-E (Nikon) using a S Plan Fluor ELWD 20x 0.45 NA objective, the microscope is equipped with a PCO.edge 4.2 bi sCMOS camera (PCO). Grid squares were automatically detected through the Fiji plugin Aveole Leonardo software using the Experimental Wizard’s, a size circular pattern to cover the grid square was micropatterned with 1000 mJ/mm^2^ of a 360 nm laser. After patterning, the grid was washed four times with DPBS. Micropatterned grids were coated with 50 ug/mL of fibronectin for 45 minutes. Approximately 25,000 RPE1 Halo-mH2A cells labeled with 1.4 nm HAN-594 were plated onto four micropatterned EM grids in a 35 mm Mattek dish (P35G-1.5-20-C). PDMS stencils 15 mm in diameter with 4 wells, each well 4 mm in diameter (Aveole #4W001) were applied to the center of the Mattek dish. An EM grid was placed within each well. Four to six hours after seeding, grids were blotted for 4-6 seconds, and then plunged into a 50/50 mixture of ethane/propane with a custom manual plunger (MPI Martinsreid) and then stored under liquid nitrogen.

### Cryo-FIB-milling and Cryo-Fluorescence

Grids clipped into FIB-compatible Autogrids (Thermo Fisher) were loaded into an Aquilos 2.0 Dual-Beam cryoFIB/SEM (Thermo Fisher). A layer of organometallic platinum was applied onto the grid. Lamellae were prepared using a Ga^2+^ ion beam at 30 kV in which the beam current was gradually reduced stepwise from 0.5 nA to 10 pA during the thinning procedure, as described in [2] and performed with Thermo Fisher AutoTEM version 2.4. The milling progress was monitored by SEM at 2 kV and 5 kV. Following automated milling, fine polishing was performed at 10 pA for 5-10 minutes. The final lamella thickness was between 140 and 180 nm. Grids were imaged by fluorescence at the Aquilos equipped with iFLM version 1.2.

### Cryo-ET data collection

#### 5 nm ribosomal dataset

Lamellae were imaged on a 300 kV Titan G2 Krios Transmission Electron Microscope (Thermo Fisher) using a K3 Summit direct electron detector equipped with a Biocontinum energy filter (Gatan) at 81,000x nominal magnification, the calibrated pixel size calibrated was 1.068 Å/pixel.

Hybrid tilt series were acquired with PACE-tomo [85]. The 0° tilt was exposed to 20 e^-^/Å^2^, while the remaining tilts were exposed to 5 e^-^/Å^2^. Tilt series range was +/-35° with a 2° increment, with a defocus range from -3 to -5 microns. Non-hybrid tilt series data tilt series range was +/-48° with a 2° increment, with a defocus range from -3 to -5 microns, total dose 180e^-^/Å^2^.

#### 1.4 nm ribosomal dataset

Lamellae were imaged on a 300 kV Titan G2 Krios Transmission Electron Microscope (Thermo Fisher) using a K3 Summit direct electron detector equipped with a Biocontinum energy filter (Gatan) at 81,000x nominal magnification, the calibrated pixel size was 1.068 Å/pixel. Tilt series were acquired with PACE-tomo [85]. The 0° tilt was exposed to 20 e^-^/Å^2^, while the remaining tilts were exposed to 4.2 e^-^/Å^2^, tilt series range was +/-35° with a 2° increment, with a defocus range from -3 to -5 microns. Non-hybrid tilt series data tilt series range was +/-48° with a 2° increment, with a defocus range from -3 to -5 microns, total dose 180e^-^/Å^2^.

#### Control ribosomal dataset

Lamellae were imaged on a 300 kV Titan G2 Krios Transmission Electron Microscope (Thermo Fisher) using a K3 Summit direct electron detector equipped with a Biocontinum energy filter (Gatan) at 81,000x nominal magnification, the calibrated pixel size was 1.068 Å/pixel. Tilt series were acquired with PACE-tomo [85]. The 0° tilt was exposed to 20 e^-^/Å^2,^ while the remaining tilts were exposed to 4.2 e^-^/Å^2^, tilt series range was +/-36° with a 2° increment, with a defocus range from -3 to -5 microns.

#### Nucleosome dataset

Lamellae were imaged on a 300 kV Titan G2 Krios Transmission Electron Microscope (Thermo Fisher) using a K3 Summit direct electron detector equipped with a Biocontinum energy filter (Gatan) at 105,000x nominal magnification, the calibrated pixel size calibrated was 0.876 Å/pixel. Tilt series were acquired, each tilt angle was exposed to 4.1 e^-^/Å^2^. Tilt series range was +/-40° with a 2° increment, with a defocus range from -3 to -5 microns.

### Data pre-processing

Individual frames were motion-corrected and gain-corrected in Warp (1.09). High dose 0° images were processed separately and then merged with the rest of the data. Frame stacks were created in Warp and aligned through a batch Etomo procedure (available on the Villa lab Github). Exposure filtering, 3D-CTF estimation, and tomographic reconstruction were performed in Warp (1.09). A custom script was used to account for the pretilt of the lamella (available on https://github.com/dgvjay/EM_Scripts:mass_normalize.py).

### Subtomogram Analysis

Ribosomal data: particles were picked on Warp generated deconvolved tomograms using cryolo version 1.8.4 [86]. Subtomograms were generated in Warp at 8 A/pixel, then imported into MATLAB version R2019b, read in by Dynamo [87]. To recover the ribosome from the 5 nm-HN labeled ribosomes through subtomogram averaging, the nanogold signal was damped, the high intensity pixel intensities one standard deviations away from the mean were replaced with randomized intensity values. Signal randomized subtomograms were used for the initial alignments, after initial centering, those angles and orientations were kept but further analysis was performed on the non-signal randomized subtomograms. For the 1.4 nm ribosomal dataset, particles were picked on Warp generated deconvolved tomograms using cryolo version 1.8.4 [86]. Subtomogram analysis was performed in Relion-3.1.4 [88] without signal randomization. For the nucleosome dataset, the tomograms were initially denoised and corrected for the missing wedge using Warp and IsoNet [89]. Subsequently, they were segmented with DeepFinder [22] to determine particle positions. Masks were applied in IMOD to exclude particles outside the chromatin region. The remaining particles were processed with a novel template-matching algorithm, specifically designed for crowded environments (Zhou et al, under review), to determine nucleosome orientations. Finally, subtomograms were extracted using Warp and further refined in Relion-4 [90] using methods in preparation.

### Visualization of 1.4 nm Nanogold in Cryo-Tomograms

Tomograms were reconstructed at 8 Å/pixel or 10 Å/pixel by WBP. Tomograms were rotated to compensate for the lamella pre-tilt using IMOD’s Rotatevol. Ten slices were averaged by means of a python script to improve the signal-to-noise (https://github.com/dgvjay/EM_Scripts/average_neighboringSlices.py). Denoised tomograms were generated using Warp’s Noise2Map program [40].

### Distance measurements

Pairwise distances were determined between refined nanogold coordinates and refined ribosome coordinates in MATLAB version R2019b. The minimum distances between each pair was plotted, and then the average and the standard deviation was determined.

### Room-temperature TEM

Cells were SLO treated as described using 4 uM 1.4 nm Alexa-594-Halonanogold and fixed in 2.5% glutaraldehyde /0.1M sodium cacodylate buffer for 5 min at RT then 2 h on ice. Cells were washed several times in 0.1M cacodylate buffer and transferred to a dark room using only a red lamp. HQ silver enhancement kit (#2012-45ML, Nanoprobes) was prepared as described and incubated with cells for 10 minutes. After this time, cells were placed on ice and washed 5 x / 0.1M cacodylate buffer. Cells were next stained with 0.7% osmium tetroxide (#19150, electron microscopy sciences) diluted in 0.1M cacodylate for 30 minutes on ice rocking. Cells were washed in water, dehydrated, and embedded in Durcupan resin (Sigma A-D) using standard protocols. 60 nm sections were obtained using a diamond knife (Diatome), and collected on formvar slot grids (VWR, FF2010-Cu-25). Images were acquired on a Biotwin TEM operating at 120 kV using a CCD camera (AMT). For ribosome quantification, images were acquired at 1900x.

### Ribosome quantification in resin embedded TEM images

Ribosome quantification was performed using Imaris v10.2.2. To quantify the number of ribosomes, a spot detection feature was applied which through thresholding, reliably detected gold particles due to their high contrast. To determine the number of ribosomes detected per cell, we first measured the average volume imaged in our TEM images, using the cell membrane to create a surface. This provides the number of ribosomes detected per cell volume. In order to extrapolate this number to the volume of the cell, the average HEK cell volume was next calculated by fluorescence imaging. Phalloidin was used to label actin filaments and thus the cell boundary, thereby allowing a surface to be created and the total HEK cell volume to be calculated. Finally, to quantify the size of gold particles, a surface feature was applied, allowing the edges of gold particles to be masked and their diameter calculated.

### ChromEMT and plastic section tomography

Cells were labeled with nanogold via SLO delivery as described. Cells were fixed in 2.5% glutaraldehyde (Ted Pella, 18420) diluted in 0.1M sodium cacodylate buffer (Ted Pella, 18851) for 5 min at RT then 2 h on ice. To allow for DNA staining, embryos were incubated with DRAQ5 (Thermo Fisher Scientific, 62251) diluted 1:500 in 0.1 M sodium cacodylate buffer for 1 h on ice. For photo-oxidation, 12mg of 3,3′-Diaminobenzidine (DAB, Sigma, D8001) was dissolved in 1 ml of 0.1M HCL by vortexing. This was diluted by adding 19 ml of 0.1M cacodylate buffer, which was passed through a 0.2µm filter. Sodium cacodylate buffer was exchanged for cold DAB solution immediately prior to photo-oxidation. This was performed using a 63x oil immersion objective and custom-built wide-field fluorescent microscope with a high-powered light source (642nm CW fiber laser, MPB Communications) and square-core multimode fiber optics (Thorlabs) to deliver the laser light into the illumination path. We experimentally determined that illuminating the sample with 100 mW of laser power for 30s-1 minute provides optimal oxidization.

### Room temperature TEM

For tomography, 300-nm sections were obtained and deposited on copper slot grids (Luxel, SKU: C-S-M-L) that were first glow discharged for 30s. For reconstruction, the grid was placed on a drop of 15 nm gold protein A gold (PAG) fiducial particles (diluted 1:50 in ddH_2_O) for 30s, before being rinsed in ddH_2_O and removing excess liquid using filter paper. These steps were repeated for the opposite side of the grid, which was subsequently left to air dry. Dual-axis tilt series were acquired on a 200 kV Tecnai F20 Transmission electron microscope (Thermo-Fisher Scientific), using an Eagle 4k x 4k CCD camera (Thermo-Fisher Scientific). The tilt series were acquired using SerialEM at a tilt range of +/-60 degrees using 1-degree increments at a magnification of 13500x, corresponding to a pixel size of 1.62 nm at a binning of 2.

### Room temperature TEM reconstruction and analysis

Dual-axis tilt series were aligned in IMOD eTomo and reconstructed with a SIRT filter. Chromatin was quantified from the final tomographic z-stacks using the MIA modularized image analysis workflow plugin (v1.2.6) for ImageJ [91].

## Extended Data

**Extended Data Figure 1.**
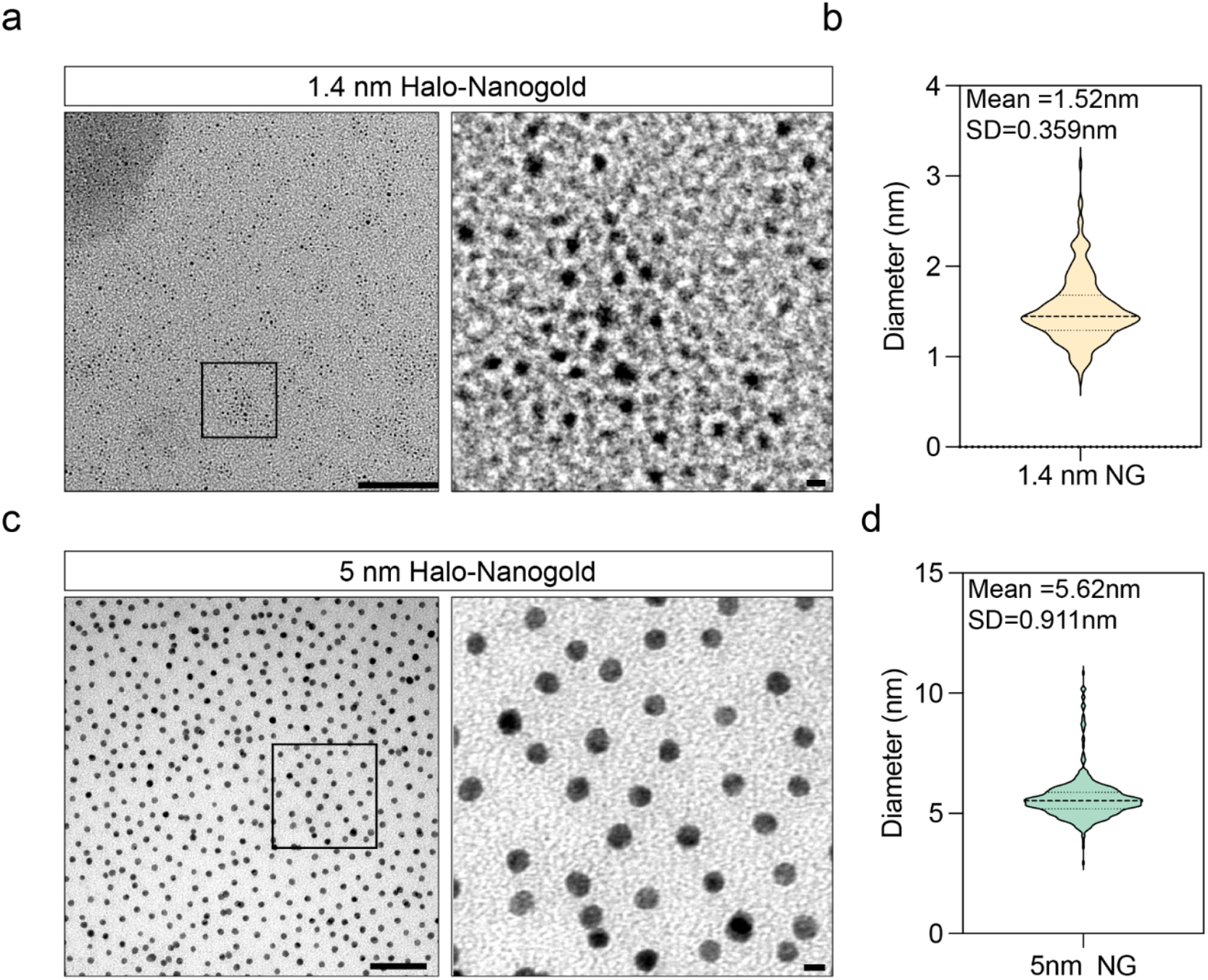
Size distribution of Halo-Nanogold particles. **(a)** TEM image of 1.4nm Halo nanogold, black inset shows zoomed region (right). Scale bar is 100 nm (left) and 1 nm (right). **(b)** Quantification of particle diameter of the image shown in a. **(c)** TEM image of 5 nm Halo nanogold, black inset shows zoomed region (right). Scale bar is 50 nm (left) and 5nm (right). **(d)** Quantification of particle diameter of the image shown in c.

**Extended Data Fig 2:**
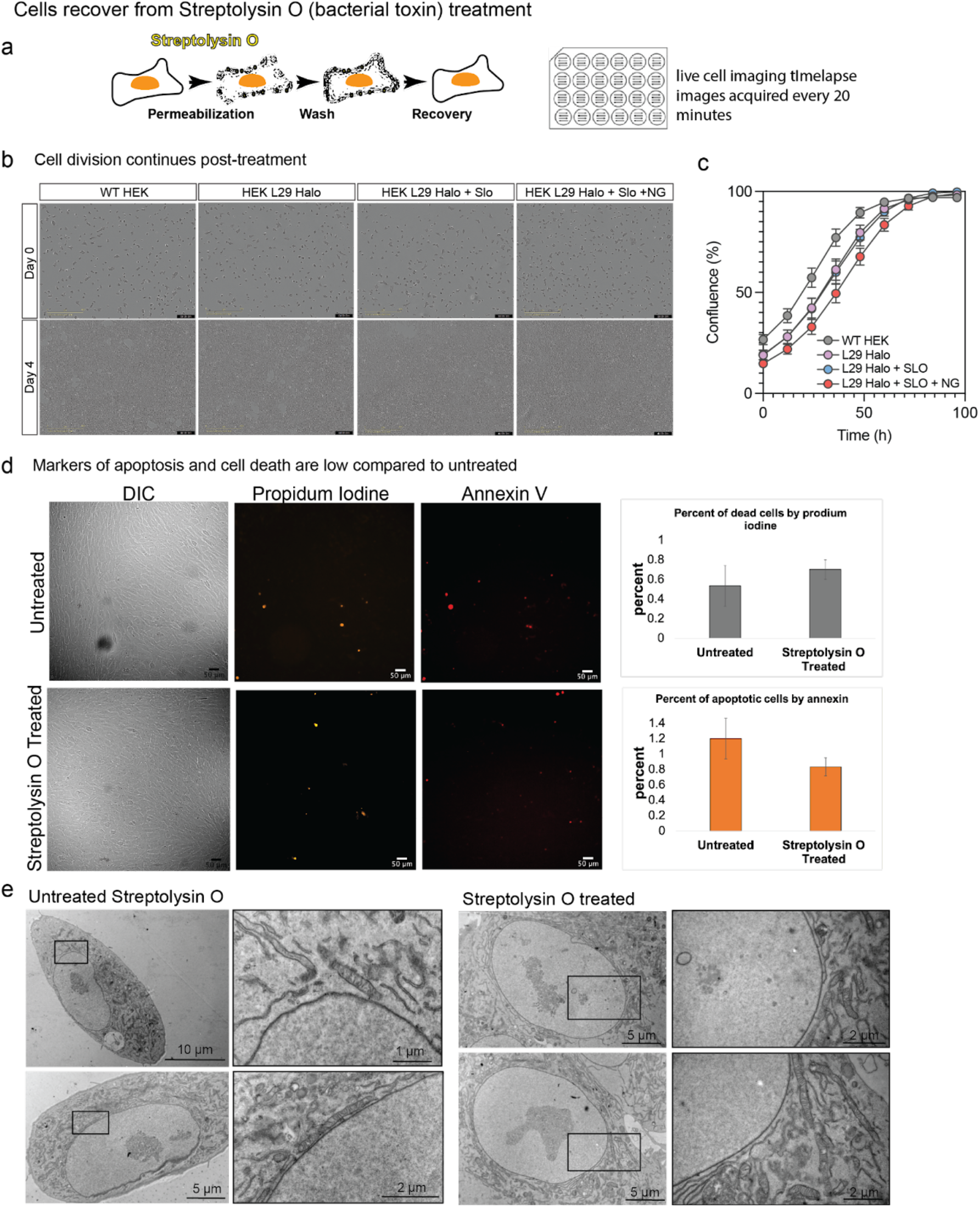
Cells recover from Streptolysin O (SLO) treatment. **(a)** Schematic of pore formation mediated by SLO treatment to mammalian cells in cell culture. (b) InCuyte images of HEK cells in the indicated conditions. Images show cell confluence at time 0 (day 1) and 100 h (day 4). Scale shows 400um. (c) graph showing the group rate of the indicated conditions. N=3 replicates, error bars show standard error. (d) Following SLO treatment, cell viability was assessed using propidium iodine (PI), and annexin V (marker of apoptosis). Quantification across three wells of cells in treated and untreated samples shows cell death and apoptosis is low. (e) The ultrastructure of the cell is maintained following SLO treatment, as seen by room temperature TEM.

**Extended Data Figure 3.**
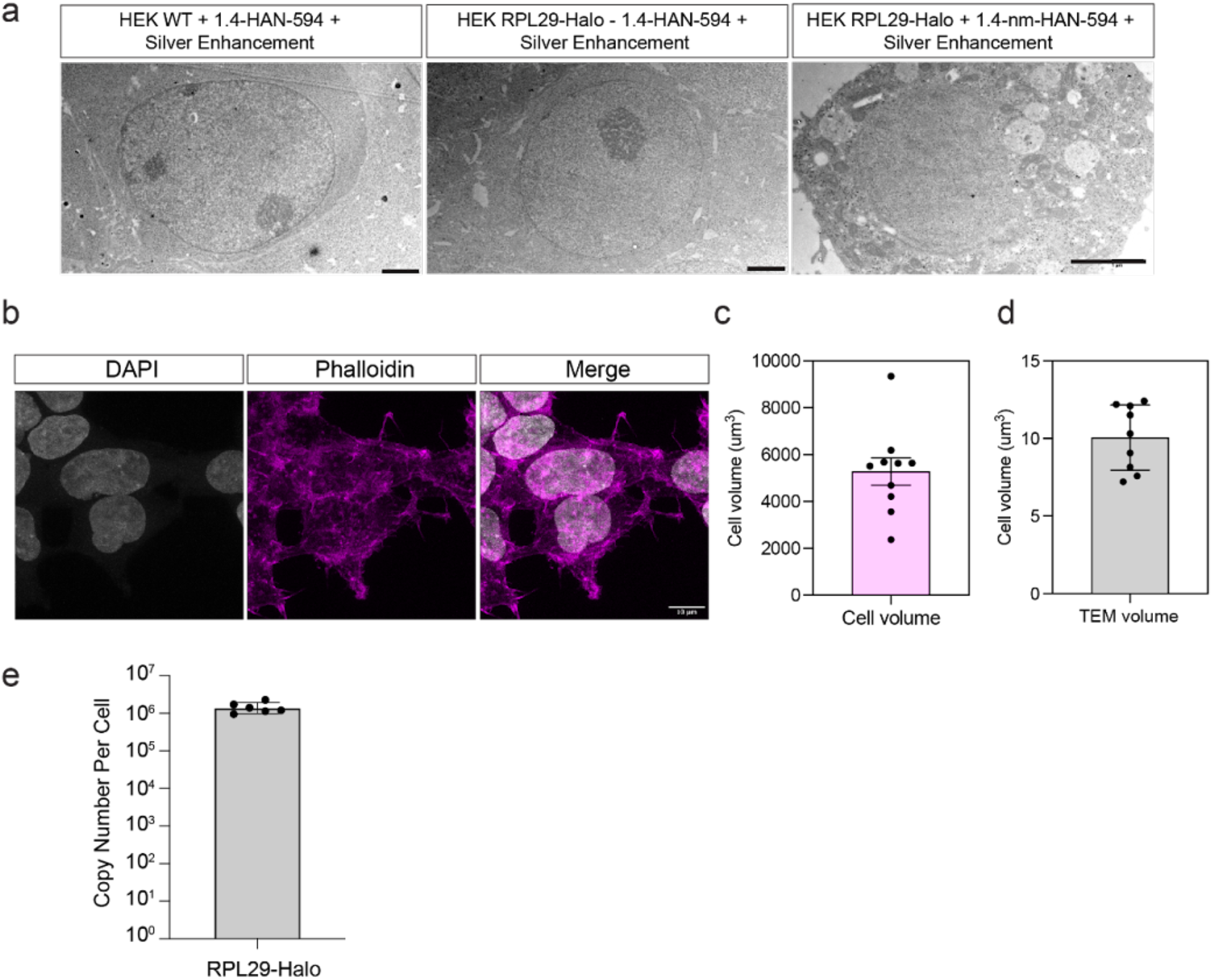
Nanogold enhancement is specific to HEK 293T RPL29-Halo. **(a)** Transmission electron microscopy (TEM) images of HEK 293 cells that were enhanced for 10 minutes. Left: WT cells without Halo-Nanogold, middle: WT cells treated with 3 uM 1.4 nm-HAN-488; right: RPL29-Halo cells treated with 3 uM 1.4 nm-HAN-488. Scale is 2um. **(b)** Immunofluorescence images of HEK 293T cells with DAPI and phalloidin staining as a maker for the cell edge. **(c)** Quantification of cell volume from the cells shown in A. N=10 cells. Mean =5283, SD=1847. **(d)** Quantification of cell volume in TEM images. N=9. Mean =10.05, SD= 2.1. **(e)** Quantification of ribosomal copy number in HEK 293 RPL29-Halo cells, N = 6 experiments.

**Extended Data Fig 4:**
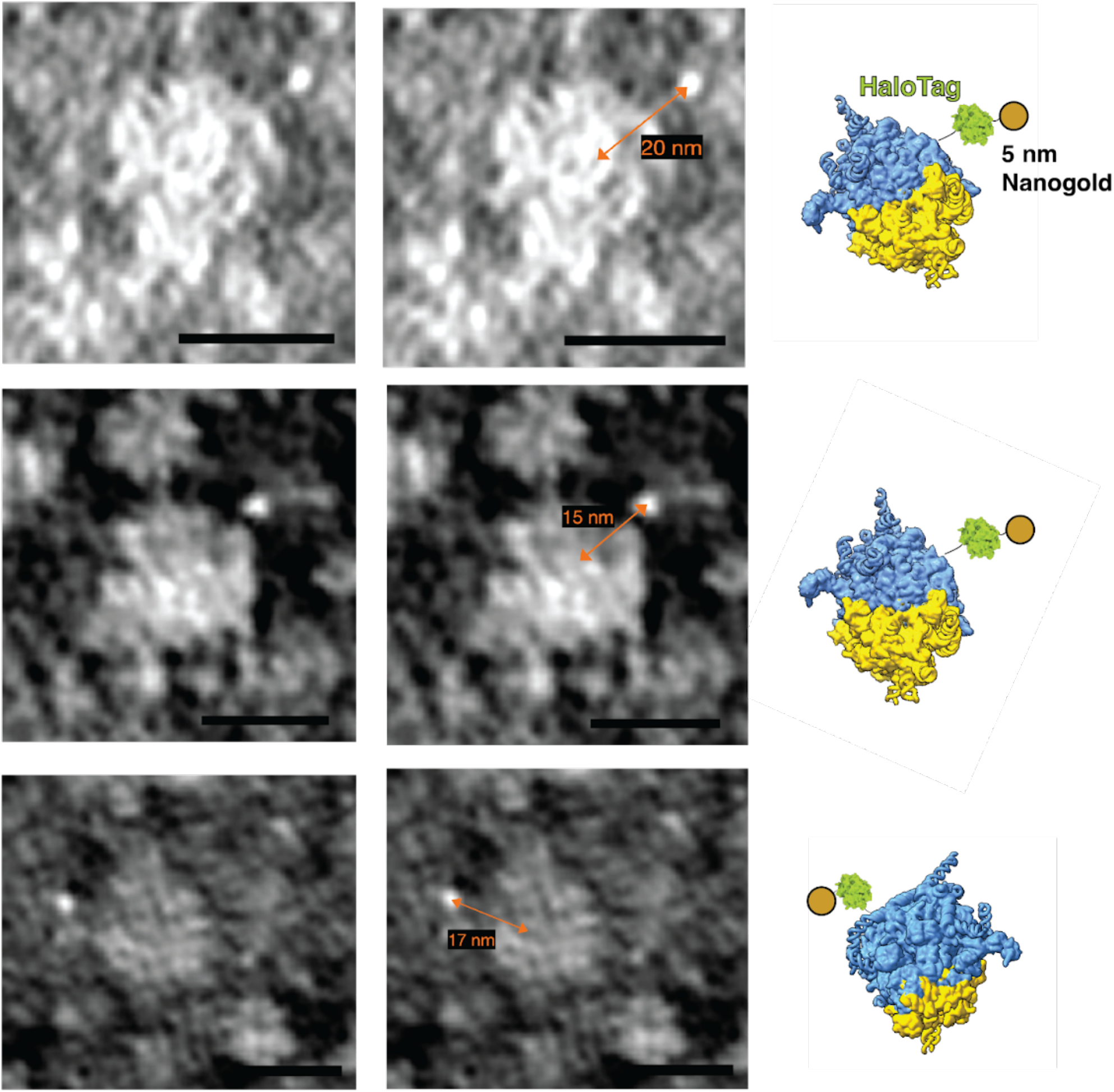
Case examples of 5 nm-HN and ribosome. Central slice of three examples of 5-nm-HN-labeled ribosomes from denoised tomograms. Contrast-inverted, i.e. white-on-black. On the right are ribosome models showing expected location of gold nanoparticle. Scale bar is 20 nm.

**Extended Data Fig 5:**
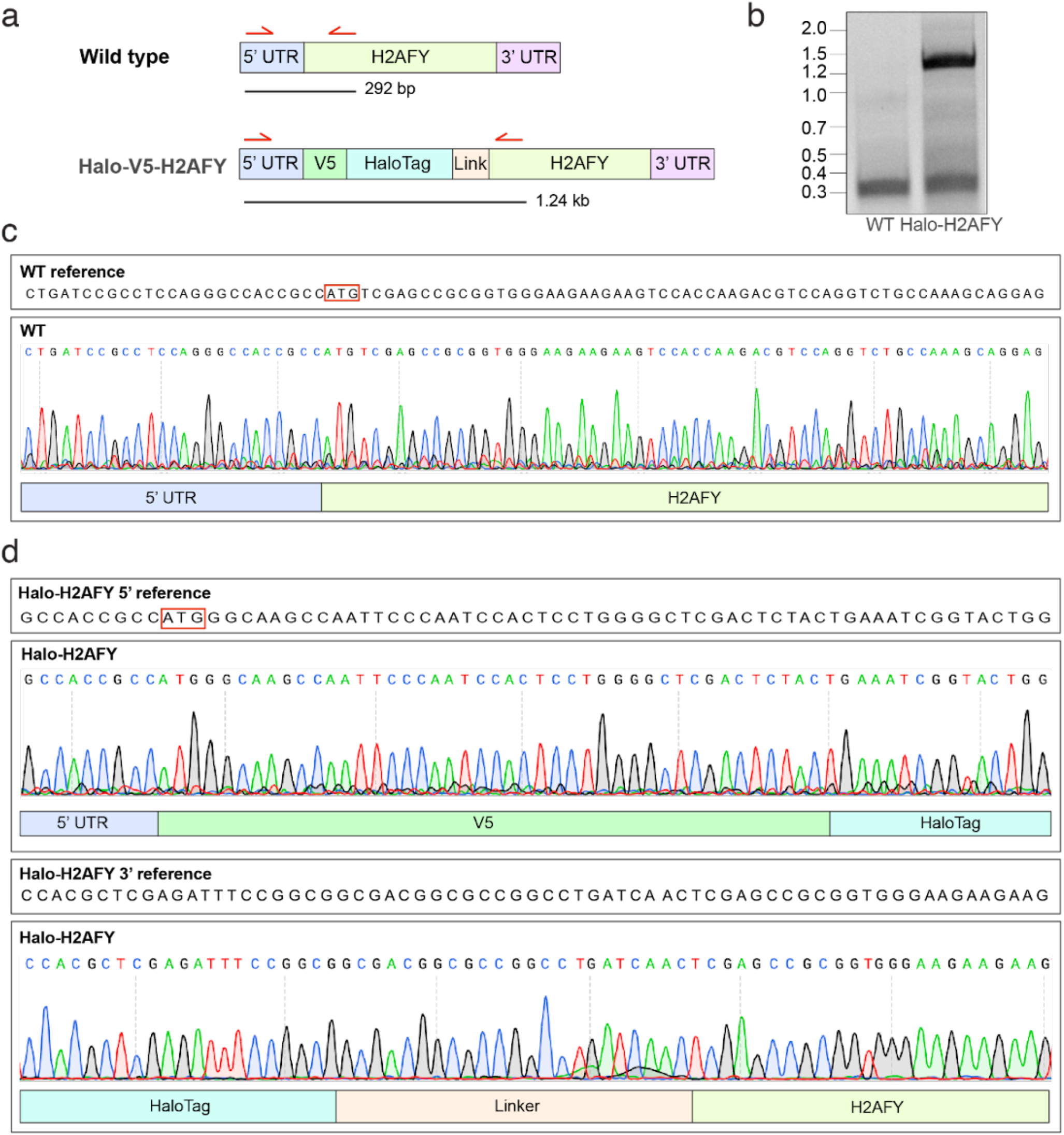
CRISPR knockin of an N-terminal HaloTag to human H2AFY. **(a)** Diagram of the knockin and PCR strategy. A V5 epitope and Halo tag are knocked into the N terminus of the human H2AFY (coding the histone variant MacroH2A) locus in retinal pigment epithelial (RPE1) cells. Top shows wild type (WT), bottom shows Halo-knockin. Red arrows highlight PCR primers used for genotyping with the expected fragment size depicted as a black bar below. **(b)** DNA gel of the PCR described in A. **(c)** Sanger sequencing of WT and V5-Halo-H2AFY RPE1 cells. **(d)** Alignment to the indicated genomic sequences.

**Extended Data Fig 6.**
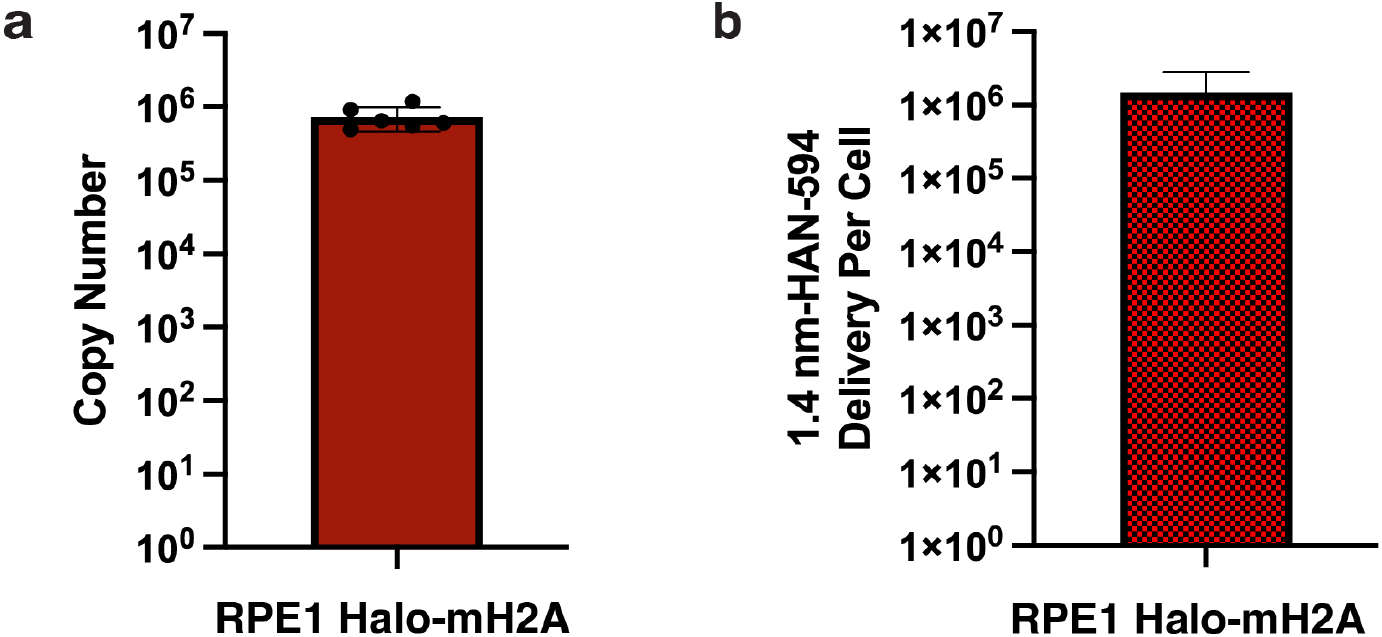
MacroH2A copy number and 1.4 nm HAN-594 delivered per cell. **(a)** Absolute copy number of macroH2A molecules per RPE1 Halo-mH2A cell. N=4 experiments. **(b)** Quantification of 1.4 nm-HAN-594 delivered to RPE1 Halo-mH2A cells, N=3 experiments.

**Extended Data Fig 7:**
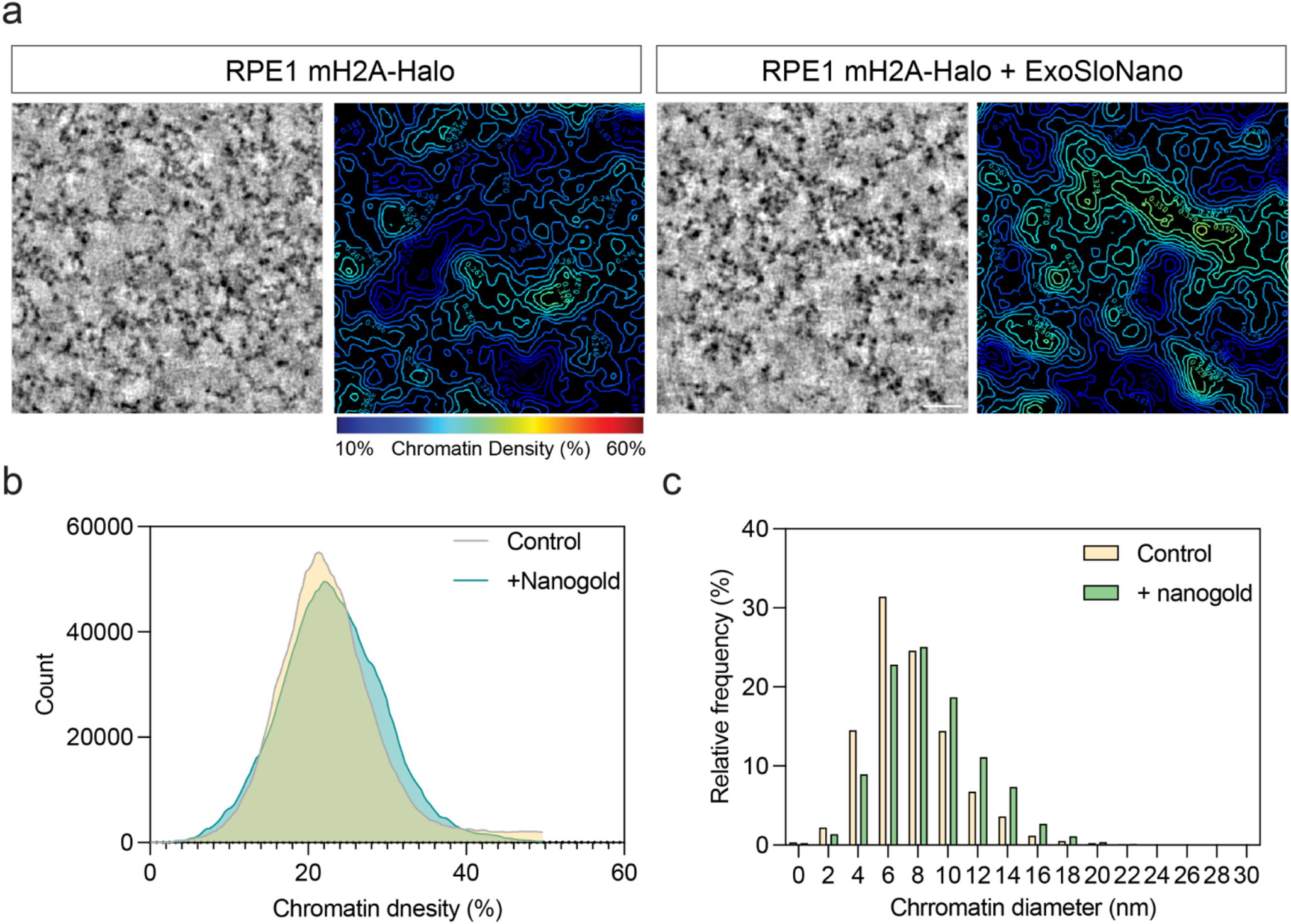
Tagging mH2A by ExoSloNano does not affect chromatin structure. **(a)** Tomographic slices of control or 1.4 nm HAN-594 treated RPE1 mH2A cells strained using ChromEMT (left). Right, shows a chromatin density map where chromatin packing is color coded based on the scale bar. Scale bar is 100 nm. **(b)** Histogram of chromatin density of the cells shown in A. N= 3 tomograms. **(c)** Histogram of chromatin diameter of the cells shown in A. N= 3 tomograms.

**Extended Data Table 1:** *In situ* cryo-FIB-ET data collection and reconstruction statistics.

|  | labeling the ribosome |  |  |  |  | labeling H2AFY |
| --- | --- | --- | --- | --- | --- | --- |
|  | in situ labeled 5 nm-HN |  | in situ labeled 1.4-nm-HAN-488 |  | in situ unlabeled | in situ labeled 1.4-nm-594 |
| Data collection |  |  |  |  |  |  |
| Nominal Magnification | 81,000x | 81,000x | 81,000x | 81,000x | 81,000x | 105,000x |
| Voltage (kV) | 300 | 300 | 300 | 300 | 300 | 300 |
| Defocus range (μm) | -3 to -5 | -3 to -5 | -3 to -5 | -3 to -5 | -3 to -5 | -3 to -5 |
| Pixel size (Å) | 1.068 | 1.068 | 1.068 | 1.068 | 1.068 | 0.876 |
| High Dose 0° | 20 | 0 | 20 | 0 | 20 | 0 |
| Total Electron exposure (e−/Å²) | 180 | 170 | 180 | 160 | 180 | 170 |
| Tilt-range/step (°) | ±36 / 2 | ±48 / 2 | ±36 / 2 | ±48 / 2 | ±36 / 2 | ±40 / 2 |
| Tilt-scheme | dose-symmetric, grouping 2 |  |  |  |  |  |
| Processing |  |  |  |  |  |  |
| Symmetry imposed | C1 |  | C1 |  | C1 | C1 |
| Initial particle images (no.) | 13,748 |  | 24,975 |  | 17,662 | 6603 |
| Final particle images (no.) | 1,743 |  | 1,100 |  | 2,7689 | 4955 |
| Map resolution (Å) | 16 |  | 10.6 |  | 11.2 | 18 Å |
| FSC threshold | 0.143 |  |  |  |  |  |

